# Mapping of DNA damage genome-wide at nucleotide resolution by circle-damage-sequencing

**DOI:** 10.1101/2020.06.28.176388

**Authors:** Seung-Gi Jin, Dean Pettinga, Jennifer Johnson, Gerd P. Pfeifer

**Author notes:** Corresponding author: Gerd P. Pfeifer, Center for Epigenetics, Van Andel Institute, Grand Rapids, MI 49525, USA.

## Abstract

To establish relationships between mutations, for example in cancer genomes, and possible mechanisms linked to DNA damage, it is necessary to know at what sequence positions of the genome the damage occurs. However, it has been challenging to specifically map DNA damage at the nucleotide level of resolution and genome-wide with high sensitivity. Here, we describe a new method, which we named circle damage sequencing (circle-damage-seq), to accomplish this goal. The method is based on circularization of DNA molecules and DNA damage-selective cleavage of the circularized DNA followed by adapter ligation and sequencing. Based on the design of this approach, only DNA damage-containing molecules are sequenced. We conducted proof-of-principle studies to show that mapping of ultraviolet B-induced cyclobutane pyrimidine dimers (CPDs) can easily be achieved and show a specific tetranucleotide sequence context for CPDs (5’PyPy<>PyT/A) with no further sequence enrichment outside of this context. Our approach shows strongly reduced levels of CPDs near transcription start sites and a spike of this damage near the transcription end sites of genes. We then show that 1,*N*^6^-etheno-deoxyadenosine DNA adducts formed after treatment of cells with the lipid peroxidation product 4-hydroxynonenal can be mapped genome-wide at adenine positions within a preferred sequence context of 5’T<u>A</u>C/G3’. The circle-damage-seq method can be adapted for a variety of DNA lesions for which specific excision enzymes are available.

## INTRODUCTION

Cancer genome sequencing has identified thousands of somatic mutations in different types of human cancer ^1–3^. These mutations provide a valuable source of information as to the potential mutagenic events that have occurred and have created the mutations, sometimes decades before the tumor develops. In some cases, convincing links between an exposure and cancer mutations have been established, best exemplified by typical sunlight-induced mutations (C to T mutations at dipyrimidine sequences) that are prevalent in melanoma and in non-melanoma skin cancers ^4^, by smoking-related G to T transversion mutations that are common in lung cancers from smokers but not nonsmokers ^5–7^, and by aristolochic acid-associated mutations in urothelial carcinomas ^8,9^. Several human cancers carry unusual and characteristic mutational signatures of completely unknown origin ^5,10,11^. Some of these signatures are thought to arise from specific DNA damage or from defective DNA repair processes, either following exposure of tissues to carcinogens or derived from endogenous chemical reactions ^9,12^.

Many DNA-reactive agents are characterized by a unique spectrum of DNA damage they produce in exposed cells or tissues that is often directly linked to the mutational spectrum. DNA damage mapping is possible at the nucleotide level within specific genes by PCR-based techniques ^13–16^. Today, it is of greater interest to develop methodology that can be used to map DNA damage at single base resolution and genome-wide. One key challenge with this type of analysis is the development of DNA damage-specific and sensitive methods.

Several such methods exist but all have certain limitations. At a lower level of resolution, antibodies against DNA lesions can be used to immunoprecipitate the damaged DNA followed by high-throughput sequencing of the collected DNA fragments ^17,18^. However, these methods do not achieve single-base resolution, as it would be required for interpreting mutational spectra linked to DNA damage.

One method is based on the release of small DNA fragments, about 20-30 nucleotides in length, by the nucleotide excision repair (NER) process within cells. These oligonucleotides can be purified from exposed cells and are then sequenced using high-throughput sequencing (excision repair sequencing, or XR-seq) ^19,20^. This method is very specific and sensitive but is limited to DNA lesions that are processed by NER and hence cannot easily detect smaller base lesions. Other methods rely on the ligation of oligonucleotides into a DNA strand break, either a single-strand break or a double-strand break ^21–26^. These methods often use secondary DNA breakage, such as by sonication, to create a random DNA break in addition to the specific DNA break produced from the base damage, which may increase background and sequencing load. Because of the limitations often associated with currently used methods, we have been developing a new approach, which we named circle-damage-sequencing (or circle-damage-seq). This method is used in combination with DNA damage-specific repair enzymes and allows the detection and mapping of DNA base damage at the single base level of resolution and genome-wide.

## RESULTS

### Overview of circle-damage-sequencing (circle-damage-seq)

DNA lesions may form as a result of radiation exposure or after exposure of cells to exogenous or endogenous reactive chemicals. In cells, the lesions can be recognized and repaired by lesion-specific DNA repair enzymes with high fidelity and efficiency. We employed these DNA repair systems to perform high-throughput mapping of various DNA lesions in damaged cells at single-nucleotide resolution on a next-generation sequencing (NGS) platform. In this approach, we adapted the circle-seq method originally based on circularization reactions to map rare genomic CRISPR/Cas9 off-target cleavage sites in vitro ^27^. We refer to this novel DNA damage-selective mapping method as “circle-damage sequencing” (circle-damage-seq) (Fig. 1).

**Figure 1:**
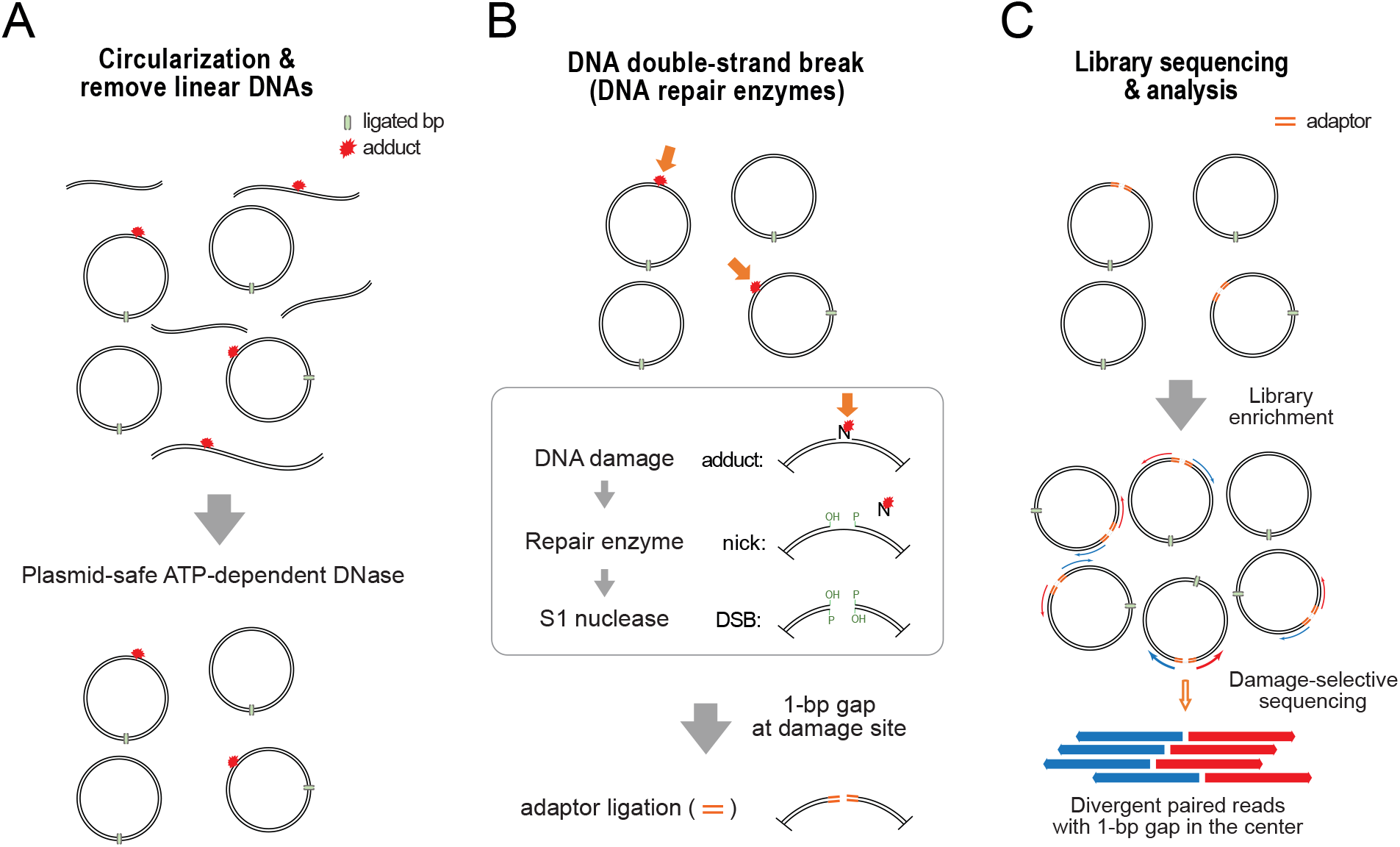
Outline of circle-damage-sequencing (circle-damage-seq). **A.** Genomic DNA is isolated from DNA damaging agent-treated cells and is sheared to obtain average size fragments of 300 base pairs. DNA is then circularized using T4 DNA ligase. Remaining non-circularized DNA fragments are removed using an exonuclease cocktail. The circular DNAs are collected after column-based purification. **B.** DNA lesions (indicated by arrows) within a DNA circle are removed by specific DNA base excision repair enzymes producing AP sites, which are then processed by the AP endonuclease, APE1, and an AP lyase. Single-strand- and nick-specific S1 nuclease cleaves the opposite strand of the nicked DNAs and leaves ligatable double-strand breaks with single base gaps. Then the sequencing adaptors are ligated to the DNA ends. Of note, in the circle-damage-seq process, non-damaged circular DNAs are not cleaved and therefore will not be sequenced. **C.** During DNA library preparation, DNA molecules opened at the circles after adduct-specific cleavage are ligated to sequencing adaptors and can be selectively enriched by PCR, representing DNA damage-specific sequence enrichment. After paired-end sequencing and alignment to the reference genome, there will be a unique aligned read pattern, divergent paired-end reads with single base gaps (corresponding to the modified base) in the center between forward (blue arrow) and reverse (red arrow) reads. Each divergent read is derived from a single opened circular DNA molecule.

We isolated genomic DNA from cells that either have an endogenous base modification or have been treated with a DNA damaging agent. In order to obtain a DNA damage spectrum after an exposure, DNA needs to be harvested before adducts can be repaired by intracellular repair activity. We treated cells with DNA damaging agents and immediately harvested the cells. Using sonication or restriction enzyme digestion, we converted the purified DNA to an average of 300 to 400 base pair size, a suitable range for DNA circularization and downstream NGS analysis. Circularization of DNA fragments smaller than 100 bp is limited. After data analysis, we obtained more uniform genomic mapping results from random shearing by sonication compared to restriction enzyme cutting, due to sequence bias of the restriction enzymes.

After the fragmentation, we performed the DNA circularization step in circle-damage-seq using 1 μg of DNA per 200 μl of reaction volume with T4 DNA ligase to minimize ligation between linear DNA fragments. As a key step in circle-damage-seq, the circularization step efficiently converts genomic DNA to a suitable size of circular DNA molecules containing modified DNA bases in random positions for next-generation sequencing. In addition to this feature, during circularization, T4 DNA ligase repairs DNA single strand breaks that are present endogenously or are caused by mechanical stress or other factors during sample preparation. Thus, the method avoids mapping most DNA single-strand breaks that are often present as background and false-positive reads in DNA adduct mapping procedures. In the next step, the remaining non-circular DNA fragments with a free end in the DNA pool are removed by treatment of the circularized DNA with an exonuclease mixture, plasmid-safe ATP-dependent DNase (Fig. 1A). Then, in order to produce double-strand breaks at DNA adduct bases within the circularized DNA, we initially convert the DNA with a lesion-specific DNA repair enzyme such as a DNA glycosylase to generate abasic (AP) sites at the positions of the damaged base. Following excision of the damaged base by the DNA glycosylase, we use AP endonuclease 1 (APE1) together with the AP lyase activity of a DNA glycosylase to cleave the AP sites, creating a one nucleotide gap in the DNA at the position of the damaged base. Next, we cleave and remove the single nucleotide on the opposite strand of the gapped DNA molecules with single-strand-specific S1 nuclease thus creating a double strand break at the DNA damage position (Fig. 1B).

As another feature of circularization, these double-strand breaks at DNA adduct positions within circularized DNAs only have one sequencing adaptor ligatable break position that can be converted into libraries suitable for next-generation sequencing. Circle-damage-seq uniquely enables sequencing of both sides of the single cleavage site in one DNA molecule using paired-end sequencing. Closed circular DNA without any DNA adduct bases is excluded from the breakage and ligation steps resulting in a lesion-selective DNA library enrichment by PCR (Fig. 1C). Notably, sequencing reads obtained from circle-damage-seq show a unique aligned read pattern, divergent reads with a single base gap in the center representing the damaged DNA base, due to the fact that each paired-end read is derived from a single opened circular DNA molecule (Fig. 1C). Thus, circle-damage-seq provides high selectivity for DNA damage mapping at single nucleotide resolution with low background and dramatically reduces the required sequencing depth and cost of this approach.

### Mapping of DpnI cleavage sites in the *E.coli* genome

We initially tested the feasibility of circle-damage-seq by analyzing the endogenous DNA modification *N6*-methyladenine (6mA). Recent work has suggested that 6mA may play important roles in gene regulation in prokaryotic and even in mammalian genomes, but 6mA is at or below the limit of detection in mammals when using currently known techniques ^28–30^. In the *E.coli* genome, 6mA is formed at 5’GATC sequences and is distributed genome-wide in dam(+) but not in dam(-) strains due to the activity of the Dam DNA methyltransferase ^31^. The DpnI restriction enzyme specifically recognizes 5’G(6mA)TC sequences with methylated adenine and cuts between G(6mA) and TC but does not cleave unmethylated 5’GATC sites (Fig. S1A). To test the applicability and sensitivity of circle-damage-seq to detect 6mA, we mixed *E.coli* DNA from two different strains, dam(+) and dam(-), to levels of 0%, 2% and 10% of dam(+) DNA in a background of dam(-) DNA representing 0%, 2% and 10% of GATC sequences being methylated at adenine in input DNA. We subjected the mixture to the circle-damage-seq procedure. DNA double-strand breakage at 6mA bases within circularized DNA was achieved through DpnI treatment which generates a blunt end by splitting the target DNA between G(6mA) and TC sequences. The converted DNA was ligated and PCR-amplified to produce sequencing libraries (Fig. S1B).

We aligned the sequencing reads to the *E.coli* K-12/MG1655 reference genome. The aligned reads were initially filtered to retain the paired reads that are divergent but have no gaps. DpnI should only cleave G(6mA)TC site and leave the beginning of the reads with 5’TC at both paired divergent reads in the sequencing output. Thus, we focused on the 5’GATC tetranucleotides context positioned at the center of divergent reads and examined the frequency of 5’GATC compared to 5’NNNN background sequences. Each library was also normalized by library size. As expected, the frequency of divergent reads with 5’GATC derived from DpnI cuts was proportional to the amount of dam(+) input DNA in the sequenced libraries (Fig. S1C). Also, the ratio of reads with 5’GATC in 2% and 10% of dam(+) libraries were significantly higher than in the 0% dam(+) control library, suggesting that circle-damage-seq is a quantitative and sensitive method to map DNA modifications in the genome. We mapped all forward reads (75 bp length) derived by circle-damage-seq to the *E.coli* reference genome and identified the enrichment and distribution of peaks along the genomic sequence. As shown at Fig. S1D, the visualized peak patterns in the IGV viewer showed the same proportional enrichment to the amount of dam(+) on the reference genome and the beginning of reads with the TC dinucleotide context, defining 6mA sites at GATC sequences.

### Circle-damage-seq for genome-wide mapping of UVB-induced cyclobutane pyrimidine dimers (CPDs) at single-base resolution

UVB-induced cyclobutane pyrimidine dimers (CPDs) at dipyrimidine sequences can be specifically recognized and cleaved by T4 endonuclease V (also known as T4-PDG) ^32^. Encouraged by the DpnI mapping results, we extended circle-damage-seq to map UVB induced CPD distribution genome-wide in mammalian cells. Mouse embryo fibroblast cells (MEFs) were irradiated with different doses of UVB. We confirmed the formation of CPDs in the genomic DNA and their effective cleavage with combined treatment of the UVB-irradiated DNA with T4-PDG and the single-strand specific S1 nuclease. This combined treatment leads to the formation of double-stranded DNA breakage at CPD sites which is easily apparent by dose-dependent DNA cleavage on non-denaturing agarose gels (Fig. S2).

To perform circle-damage-seq, we fragmented the isolated genomic DNAs from UVB treated cells to an average of 300 bp size of DNA by using sonication and then performed the circularization step. As shown in Fig. 2A, to create double-strand breaks at the DNA lesions, we initially used the CPD-specific DNA glycosylase, T4-PDG, which has AP lyase activity, and the AP endonuclease APE1, to cleave the 5’ side of the sugarphosphate backbone at pyrimidine dimers. We followed this treatment by incubation with *E.coli* photolyase under long-wave UVA light to remove dimerized pyrimidine bases that remain after the DNA glycosylase incision. Next, S1 nuclease treatment generated ligatable double-strand breaks within the circularized DNA. Note that the combined treatment creates a single nucleotide gap representing the 5’ pyrimidine of the dimer.

**Figure 2:**
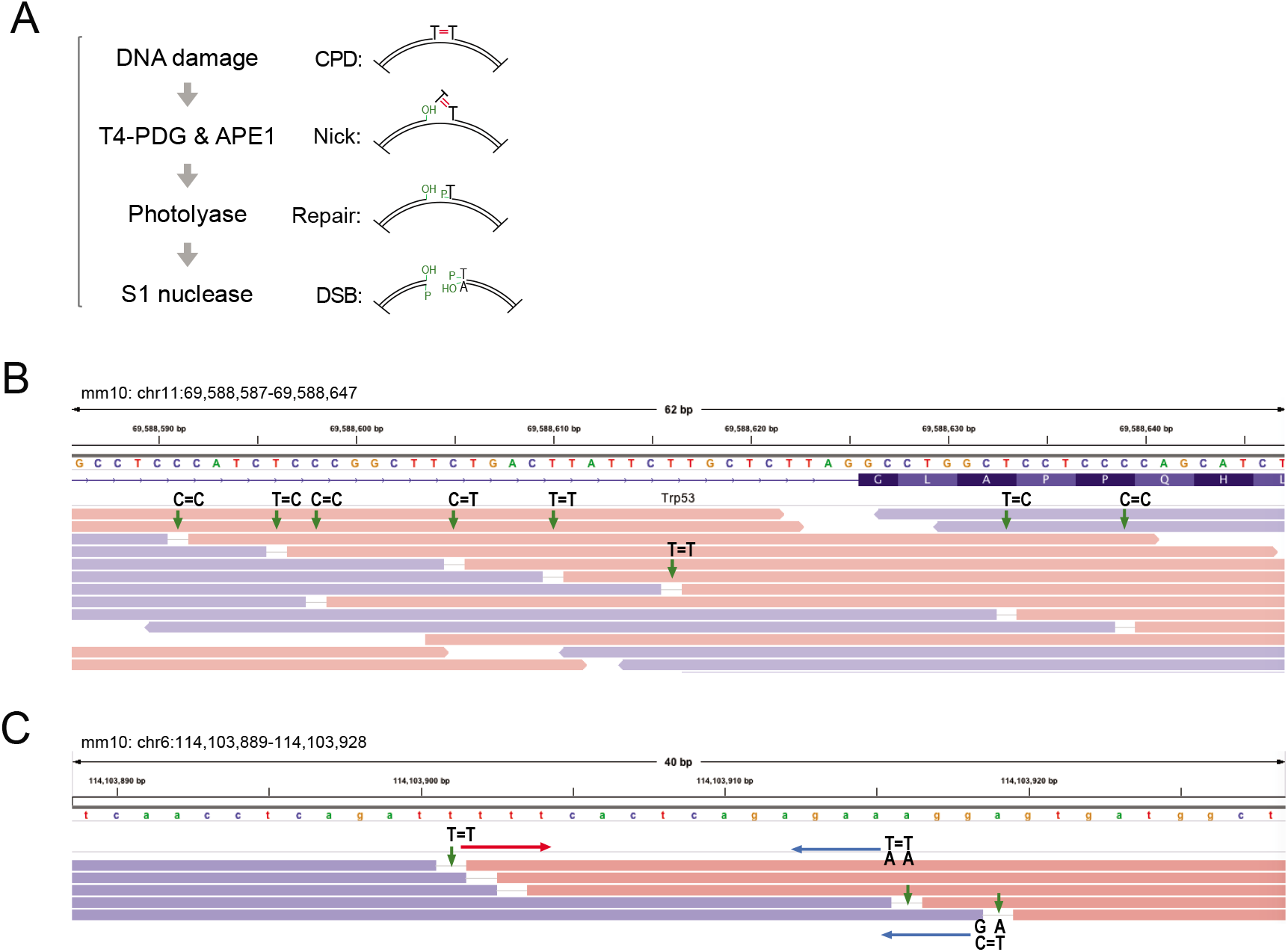
Genome-wide mapping of CPD lesions and visualization at single nucleotide resolution. **A.** CPD lesions at dipyrimidine sequences were induced into the MEF genome by UVB irradiation. To create double-strand breaks at CPDs (e.g., T=T; thymine dimer, red), CPD specific-DNA glycosylase (T4-PDG) and APE1 coupled enzyme reactions were initially used and followed by *E.coli* photolyase treatment to create ligatable, nicked DNA. To create double strand breaks, S1 nuclease was used to generate a free 5’ phosphate and 3’ hydroxyl group at cleaved DNA ends, which are ligated and sequenced. **B.** Snapshot of the IGV viewer displays the characteristic divergent paired reads of CPD mapping by circle-damage-seq aligned to the mm10 mouse reference genome at singlebase resolution (chr11:69,588,587-69,588,647). Red segments represent reads mapped to the plus strand (forward) and blue segments represent reads mapped to the plus strand (reverse). Each paired-end read is derived from a single DNA circle and is displayed as a single divergent pair. Green arrows indicate single-base gaps matching pyrimidine bases on the reference sequence corresponding to the 5’ base of a pyrimidine dimer. **C.** Snapshot of the IGV viewer represents CPD sites mapped by circle-damage-seq at an aligned region (chr6:114,103,889-114,103,928) at single-base resolution.

We aligned the sequencing reads from circle-damage-seq to the mm10 mouse reference genome. To evaluate the divergent read patterns, we converted the mapped reads obtained from 1,000 J/m^2^ of UVB-irradiated MEF cells to the bam file format and visualized the reads at single-base resolution using the IGV viewer. Indeed, we observed the divergent paired reads with a single base gap in the center and confirmed the matching bases in the gaps as pyrimidines on the reference sequence that have another pyrimidine base right next to it in the 3’ position. We observed this pattern with a high frequency at 5’TT and 5’TC sites and at lower frequency at 5’CT and 5’CC dinucleotide sequences. This result is consistent with the known specificity of CPD formation at dipyrimidine sequences (Fig. 2B and Fig. S3A) ^4^. We also observed a dipyrimidine sequence context on the opposite DNA strands of the reference genome, confirming that circle-damage-seq can detect DNA damage on either DNA strand (Fig. 2C). These divergent reads were distributed relatively uniform across the genome and can be displayed at the gene resolution level as shown, for example, in Figure 2 and in Figure S3B.

Previous work has focused mainly on the dinucleotide specificity of genomic UV damage ^4,26,33–35^. First, we confirmed that CPDs were most frequent at 5’TT and 5’TC dipyrimidines followed by 5’CC and 5’CT. We next analyzed trinucleotide sequences that reflect the sequence-specificity of CPD formation using approximately 200 million aligned, divergent reads with single nucleotide gaps. The most frequent sequence contexts considering the base flanking the dimerized pyrimidines at the 3’ side were 5’TTT and 5’TTA and these were followed by 5’TCT and 5’TCA (Fig. 3A). Thus, T or A are the preferred bases 3’ to the most abundant CPDs (5’TT and 5’TC). The most highly enriched trinucleotide sequences considering the 5’ neighboring base of the CPDs were 5’TTT and 5’CTT, followed by 5’CTC and 5’TTC (Fig. 3B) showing a preference for having another pyrimidine 5’ to a pyrimidine dimer. Divergent reads with single base gaps were >300 times less frequent in non-treated (NT) cells than in UVB-irradiated cells (Fig. 3). There was no particular enrichment of sequencing reads associated with such trinucleotide sequences in the non-UVB-treated control samples (Fig. 3A, 3B, bottom panels).

We then analyzed a potential tetranucleotide sequence context for CPD formation (Fig. S4). For the two bases 5’ of the CPDs (5’Py<>PyNN), we observed the most frequent contexts as 5’TTTT, 5’ATTT and 5’CTTT (Fig. S4A). For the two bases 3’ of the CPDs (5’Py<>PyNN), we observed the most frequent contexts as 5’TTTA, 5’TTAA and 5’TTTT (Fig. S4B), again reflecting a preference of T or A as the base neighboring a CPD at the 3’ side. It should be noted that 5’CG sequences are 5-6-fold underrepresented in mammalian genomes and this underrepresentation has not been corrected in the current analysis.

**Figure 3:**
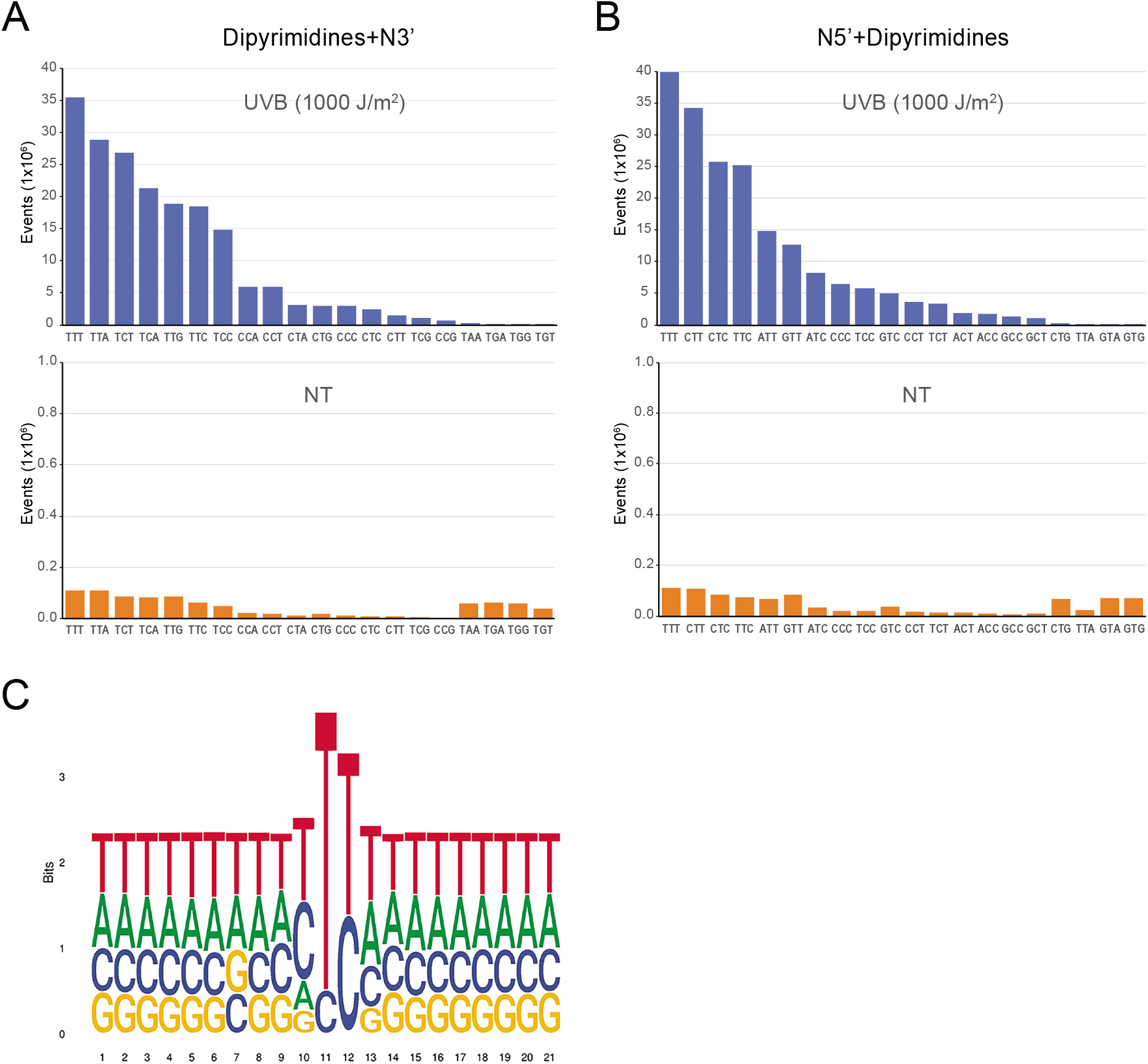
CPD mapping by circle-damage-seq shows CPD sequence specificity. **A.** Distribution plot of trinucleotide sequences (5’YYN3’) undergoing CPD formation in UVB-irradiated cells (blue). NT, non-treated control (orange). **B.** Distribution plot of trinucleotide sequences (5’NYY3’) undergoing CPD formation in UVB-irradiated cells (blue). NT, non-treated control (orange). **C.** Logo plot shows the accumulated nucleotide composition relative to mapped CPD positions along the reference genome. Position 11 represents the nucleotide in the gap between divergent reads, and position 11 and 12 represent UVB-damaged dipyrimidine sequences undergoing CPD formation. The total height of the stack represents the conservation of that position (measured in bits) using Shannon entropy. Within each stack, the height of each letter represents the relative frequency of that nucleic acid base.

To identify possible broader sequence motifs for CPD formation, we obtained a base enrichment on twenty nucleotide regions around the gapped base-pair (Fig. 3C). Position 11 represents the nucleotide in the gap between divergent reads (the 5’ pyrimidine of the CPD). We observed a dominant enrichment of 5’TT or 5’TC contexts at positions 11 and 12 representing the pyrimidine dimers. In this motif, T or C were enriched in position 10 and T and A bases were enriched in position 13. Surprisingly, however, no further sequence enrichment was observed outside of this tetranucleotide context (positions 10 to 13). Our findings further define the enrichment of specific trinucleotide and tetranucleotide sequences as most susceptible to CPD formation in UVB-irradiated cells and define a new tetranucleotide sequence preference for this type of UV damage. This data confirms the high sensitivity and specificity of circle-damage-seq.

We performed sequencing of up to 0.8 billion paired-end reads from a single circle-damage-seq library from cells irradiated with 1000 J/m^2^ of UVB. Figure 4 shows increasing read density as a function of the number of reads obtained. Next, we examined the global distribution of CPDs detected by circle-damage-seq in the mouse genome. We observed that CPD formation in cells is chiefly uniform along the genome and is largely dependent on DNA sequence where 5’TY sequences show the strongest accumulation of CPDs.

**Figure 4:**
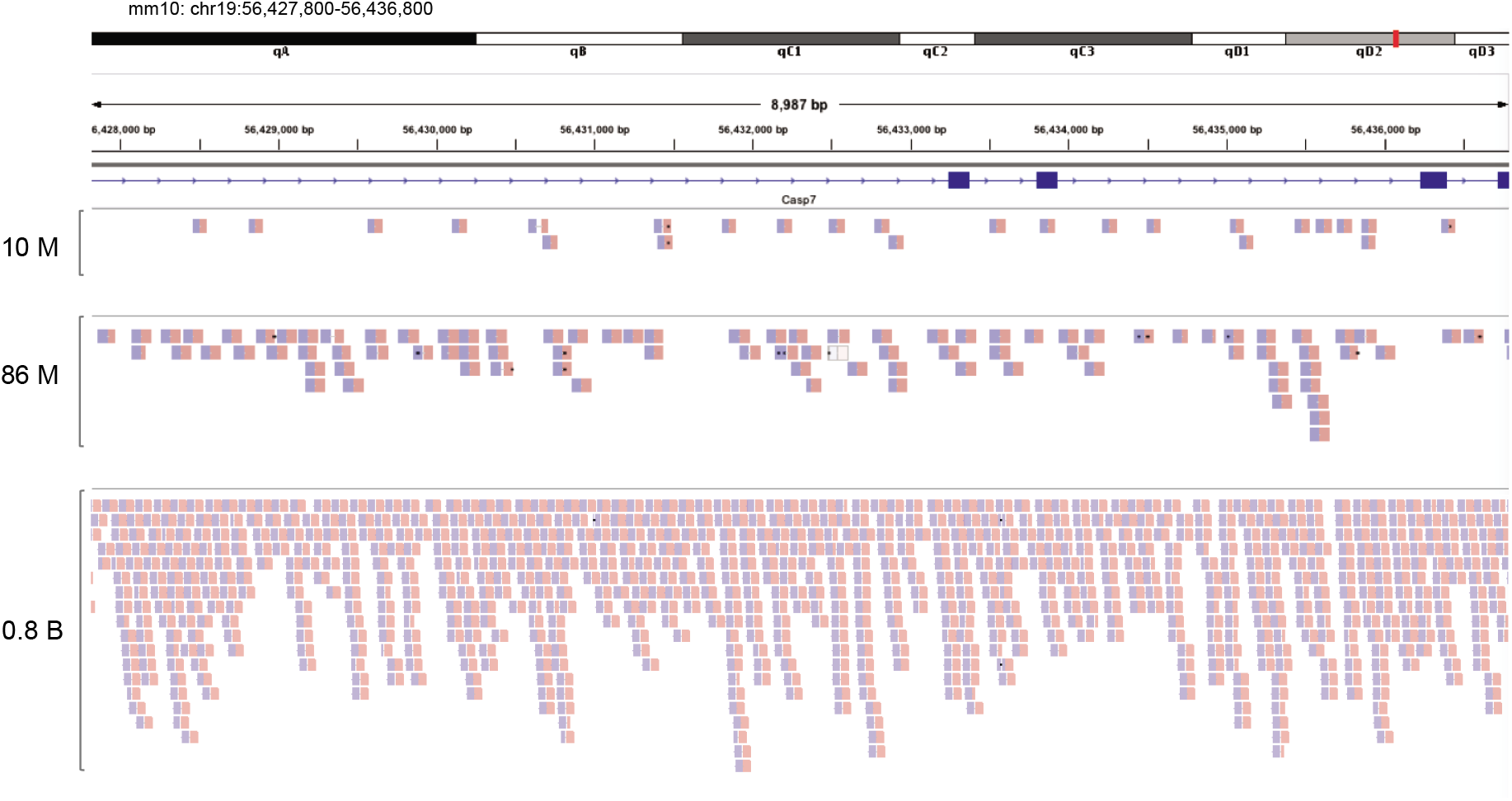
Coverage comparison of CPD lesions relative to sequencing depth. The snapshot of the IGV viewer shows the read density maps for 9 kb windows (chr19:56,427,800-56,436,800) of the mm10 mouse reference genome for the *Casp7* gene. Each read depth was obtained from circle-damage-seq libraries prepared with 1000 J/m^2^ of UVB dose-irradiated cells. The divergent, paired reads of CPD mapping by circle-damage-seq were displayed as paired blue and red segments. 10 M; 10 million reads, 86 M; 86 million reads, and 0.8 B; 0.8 billion reads.

Interestingly, we observed low levels of CPD formation at many gene promoter regions (Fig. 5 and Fig. S5). Promoter regions are often G+C-rich and provide less opportunity for T-containing CPDs to form. Indeed, plotting of the G+C content of genes shows a pronounced increase near transcription start sites, which is reverse to the CPD plot (Fig. 5B, 5C). Examples of specific genes are shown in Figure 5D and in Figure S5. In addition to sequence-based reduction of CPDs at promoters, this finding is also consistent with previous observations reported a number of years ago for single gene promoters for which we showed that transcription factors most often repress CPD formation at many sites within promoter regions ^14,36^, and as also seen in more recent reports ^25,26,35,37,38^.

**Figure 5:**
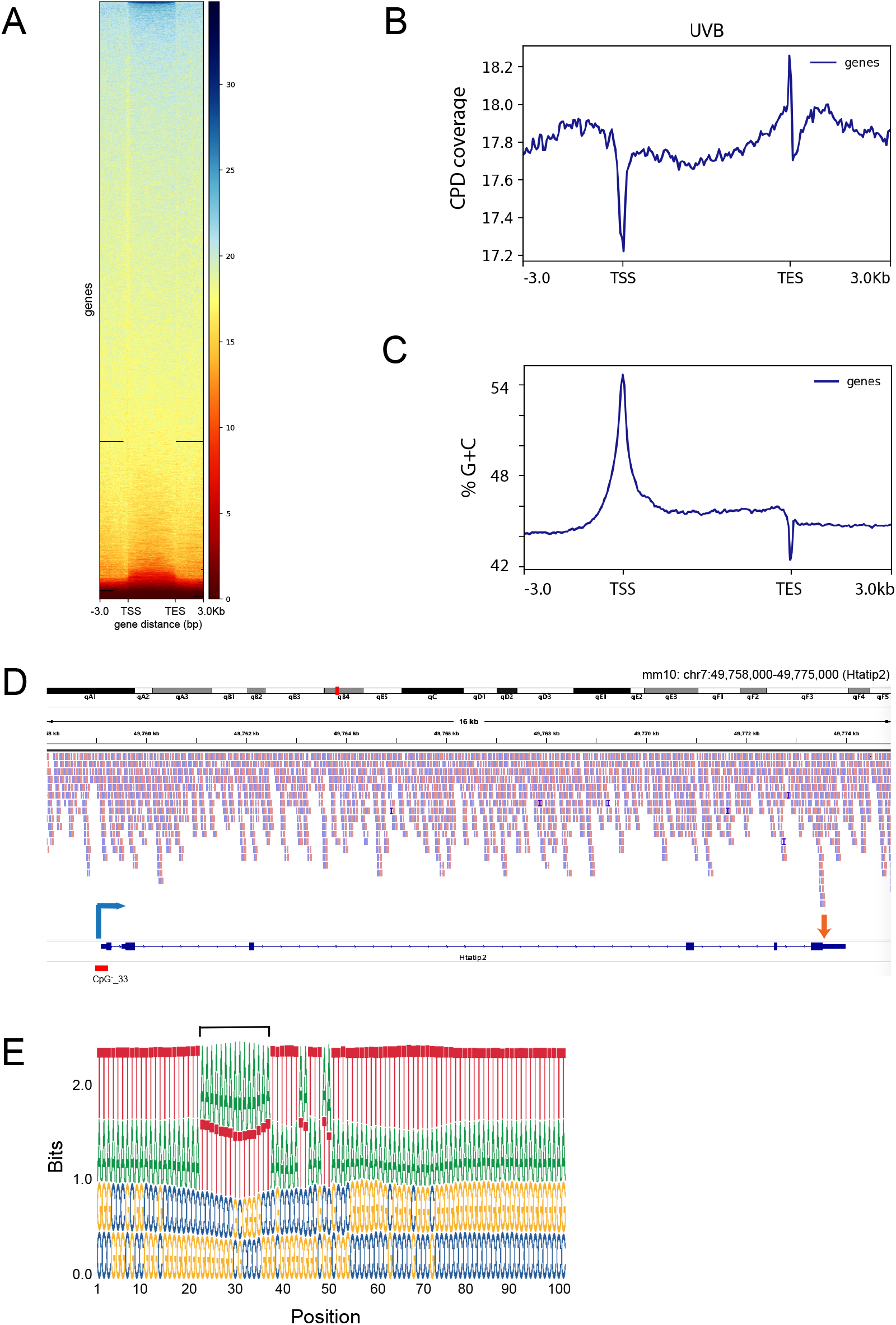
Gene-level distribution of UVB-induced CPDs shows reduced levels near the transcription start sites and a spike of CPDs near the transcription end sites. **A.** Heat map of CPD distribution along all genes of the mm10 mouse genome. CPD coverage is sorted from high (top) to low (bottom). The CPD signals were mapped and binned in 50 bp windows from 3 kb upstream of the TSS, then normalized relative to gene length over the gene bodes to the TES, and 3 kb downstream of the TES. **B.** Profile of CPD distribution along all genes. **C.** Profile of the G+C content (%) along all genes (50 bp windows). **D.** Example of a typical CPD sequence read distribution along the *Htatip2* gene. This gene shows a reduced frequency of CPDs near the TSS (blue arrow) and a spike near the TES (red arrow). **E.** Logo plot of sequences near the transcription end sites of all mouse genes (+/- 50 nucleotides surrounding the transcription end sites (TES). Note the general A/T-richness and the pronounced enhancement of A/T near the polyadenylation signals 10 to 30 nucleotides upstream of the actual transcript end, which is at position 50.

The meta-gene profile of CPD formation along all genes in the mouse genome not only shows an overall dip of CPDs near the transcription start sites (TSS) but reveals a striking spike of CPD formation near the transcription end sites (TES) of genes (Fig. 5; Fig. S6). It is likely that the enhanced susceptibility of these regions to CPD formation is related to DNA sequence features near the TES. A preferred TT-containing CPD sequence context is inherent to human and mouse polyadenylation signals. The two preferred sequences are 5’AAUAAA, which is present in almost 60% of mouse genes and the second most preferred one is 5’AUUAAA, which is found in ~16% of mouse genes ^39^ (Fig. 5E). Sequences surrounding these motifs are additionally T/U-rich and include Urich or U/G-rich elements that are located 20–40 nt downstream of the cleavage site. There is also widespread alternative polyadenylation in human and mouse genes ^39^.

### Genome-wide mapping of etheno-dA adducts using the circle-damage-seq method

To demonstrate that circle-damage-seq can be applied to a variety of DNA adducts, we mapped the distribution of 1,*N*^6^-etheno-deoxyadenosine (etheno-dA) adducts in 4-hydroxynonenal (4-HNE) treated human liver cells. As one of the major lipid peroxidation products, the 4-HNE aldehyde forms exocyclic etheno-dA DNA adducts, which are known to have mutagenic and carcinogenic properties ^40–42^. These DNA adducts are detectable in human liver using sensitive mass spectrometry-based methods and have therefore been linked to liver carcinogenesis ^43^. However, no methods are currently available to map etheno-dA adduct positions in the genome at single-nucleotide resolution.

We treated an immortalized human hepatocyte cell line, THLE-2, with 40 μM 4-HNE for 10 minutes to induce etheno-dA DNA adducts. We isolated the genomic DNA and fragmented it using NlaIII restriction enzyme, which cleaves at 5’CATG sequences and leaves a sticky DNA end to enable effective circularization of DNA fragments. As shown in Figure 6A, to create double-strand breaks at these DNA lesions, we used the monofunctional DNA glycosylase, hAAG that cleaves the *N*-glycosidic bond of etheno-dA ^44^. After hAAG incision, we used APE1 plus an AP lyase to create the adduct-specific nucleotide gaps and followed this treatment with S1 nuclease digestion to generate a ligatable double-strand break at the etheno-dA sites within the circularized DNA. The break points were labeled with an Illumina sequencing adaptor, and the library was sequenced with paired-end reads to obtain about 15 million reads.

**Figure 6:**
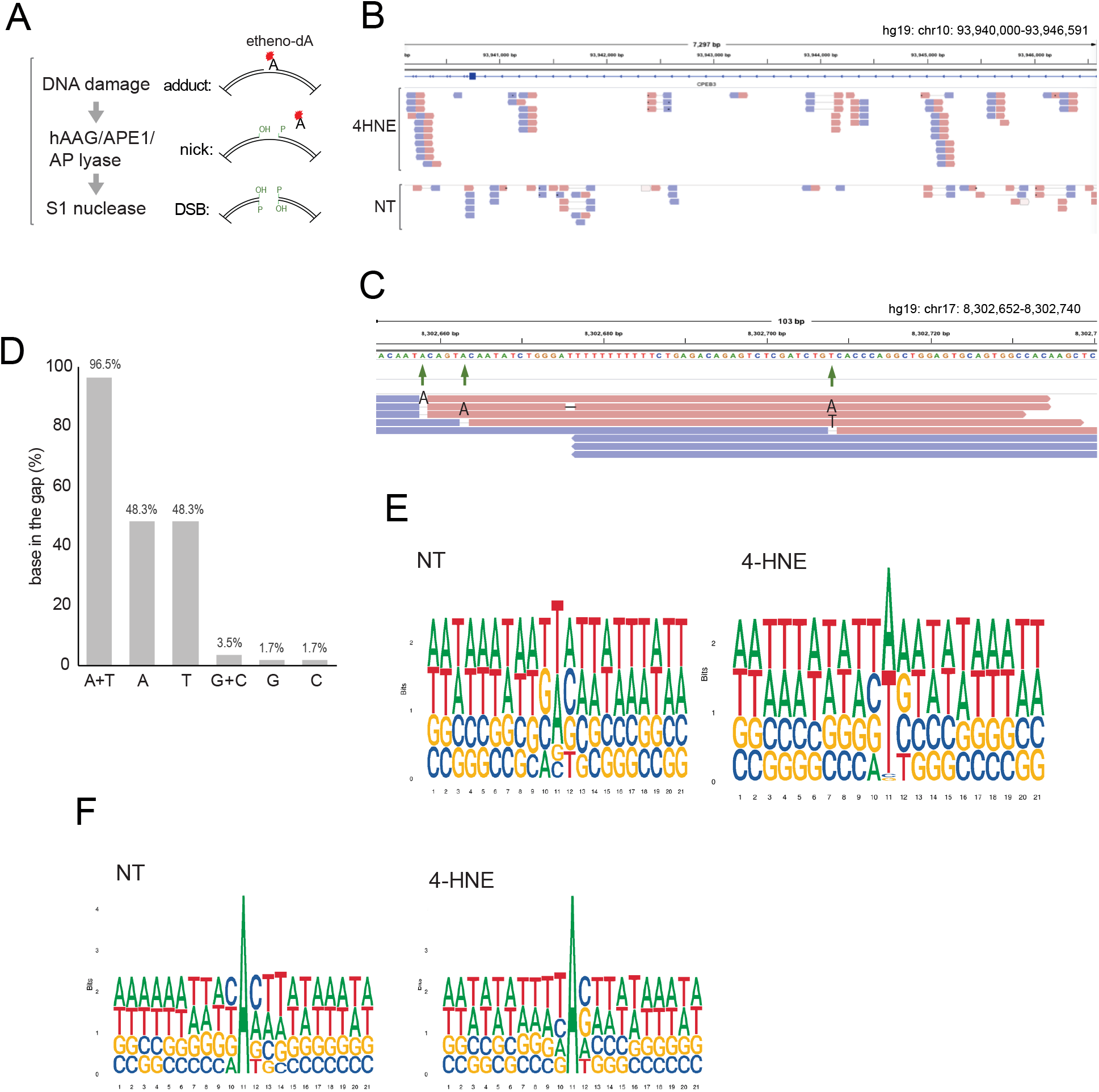
Genome-wide mapping of etheno-dA adducts using circle-damage-seq. **A.** Etheno-dA adducts were induced in the THLE-2 genome by treatment with 40 μM of 4-HNE. To create double-strand breaks at etheno-dA (A with red symbol), the etheno-dA specific-DNA glycosylase hAAG and APE1/AP lyase coupled enzyme reactions were used to create nicked DNA. Then, S1 nuclease was used to generate a free 5’ phosphate and 3’ hydroxyl group at cleaved DNA ends, which are ligated and sequenced. **B.** The snapshot of the IGV viewer displays the divergent, paired reads of etheno-dA mapping by circle-damage-seq. The distribution map shows 6.6 kb windows (chr10:93,940,000-93,946,591) of the hg19 human reference genome. Each paired-end read derived from a single DNA molecule was displayed as a single divergent pair (paired blue and red segments). 4-HNE; 4-HNE treated THLE-2 sample. NT; non-treated control THLE-2 sample showing almost no paired, divergent reads with single base gaps. **C.** The snapshot of the IGV viewer represents etheno-dA sites mapped by circle-damage-seq at single-base resolution (chr17:8,302,652-8,302,754). Red segments represent reads mapped to the plus strand (forward) and blue segments represent reads mapped to the plus strand (reverse). Green arrows indicate single-base gaps matching etheno-dA bases in the reference sequence. **D.** Percentages of nucleotides positioned in the gap between divergent reads. ‘G + C’ and ‘A + T’ indicate the combined percentages of G and C or A and T in the gaps, respectively. **E.** Logo plot bit scores of DNA bases modified in 4-HNE-treated cells and in non-treated (NT) controls. This analysis considers both DNA strands. **F.** Logo plot bit scores of modified adenine bases in 4-HNE-treated cells and in nontreated (NT) controls. This analysis is specific for adenine bases in the gapped positions and reflects the single strand-specific DNA sequence context.

The paired sequencing reads were aligned to the hg19 human reference genome. We observed evenly distributed divergent paired reads genome-wide from the 4-HNE treated liver cell samples, but we hardly found any divergent reads from control cell samples not treated with 4-HNE (Fig. 6B). In addition to obtaining the expected divergent read pattern, our bioinformatics analysis showed that the vast majority (96.5%) of the reference DNA bases matching to the single base gaps between divergent reads were adenines or thymines (depending on the targeted DNA strand) (Fig. 6C and 6D), as expected from etheno-dA DNA adducts. This data further confirms that circle-damage-seq is a highly sensitive technique applicable for genome-wide mapping of various DNA lesions at single-nucleotide resolution whenever a suitable DNA repair enzyme is available to specifically excise the lesion.

We then analyzed the sequence context in which the etheno-dA adducts are formed. The logo plots confirm the prevalence of A or T as the gapped nucleotide (Fig. 6). This base pair is also enriched moderately in the non-treated (NT) sample, although there were very few such gapped, divergent reads. Potentially, this low background may arise from rare endogenous 3-methyladenine bases, which are also excised by AAG DNA glycosylase ^45^. Examining the 5’ and 3’ nearest neighbor bases in the single-strand sequence context of adenines, we noticed an enrichment of T as the 5’ base and of C or G as the 3’ base, which suggests that 5’T<u>A</u>C/G is a preferred sequence context for etheno-dA DNA adduct formation in human cells.

## DISCUSSlON

We describe a new method that can be used to analyze modified DNA bases in mammalian cells at the single nucleotide level of resolution and genome-wide. The method depends on the ability to convert the modified base into a DNA single-strand break. Cleavage with S1 nuclease on the opposite DNA strand creates an adduct-specific ligatable double strand break. DNA glycosylase enzymes which operate at the initial step of the base excision repair pathway ^46^ can easily be adapted for this technique. Monofunctional DNA glycosylase cleave the N-glycosidic bond of the modified base and leave behind an AP site, which is further cleaved by AP endonuclease in cells. Bifunctional DNA glycosylases additionally cleave the AP site through their AP lyase activity after removal of the modified base ^47^. The combined cleavage of DNA lesions with DNA glycosylases, APE1, AP lyase and S1 nuclease creates ligatable ends and a single nucleotide gap, which is then assigned to the modified base after genome sequencing. A variety of DNA glycosylases may be employed for circle-damage-seq. Pyrimidine dimer specific DNA glycosylase and AAG protein were employed here to map CPDs and etheno-dA adducts, respectively. The oxidized guanine-specific enzyme OGG1 could be used for mapping of 8-oxo-deoxyguanosine in DNA (8-oxo-dG). DNA glycosylases of the NEIL family, e.g. NEIL1, may also be used for mapping of 8-oxo-dG. These enzymes additionally recognize other oxidized guanine derivatives including spiroiminodihydantoin and guanidinohydantoin ^48,49^. Oxidized pyrimidine bases may be excised by NTH family proteins. Uracil arising in DNA from cytosine deamination could be excised and mapped using uracil DNA glycosylases.

It should also be possible to adopt the nucleotide excision repair (NER) complex for mapping of DNA adducts, in particular bulky adducts, which are excellent substrates for this repair pathway. When the NER system of *E. coli* (UvrABC complex) is used, the incisions on the damaged DNA occur, dependent on the exact lesion to be processed, approximately seven bases 5’ to the lesion and four bases 3’ to the lesion ^50,51^. In some cases, for example for excision of benzo[*a*]pyrene-derived N2-guanine adducts, the positions of the excision are well defined ^52^, which should permit base-resolution mapping of these DNA adducts. The longer single-stranded gap remaining after excision with NER enzymes and removal of the damaged fragment should be a good substrate for S1 nuclease leading to an 11-nucleotide gap of each divergent read after circle-damage-seq.

Modified DNA bases not only may arise from DNA damage but they can represent endogenous modifications with an epigenetic function. Examples are the *N*^6^-methyladenine bases, which we have mapped here at DpnI sites (Fig. S1). Other important modified bases are derived from cytosine or from 5-methylcytosine (5mC) and include a series of oxidized 5mC bases in the form of 5-hydroxymethylcytosine (5hmC), 5-formylcytosine (5fC) and 5-carboxylcytosine (5caC) ^53,54^. There are restriction enzymes that can recognize and distinguish these modified cytosine bases and could be used for their mapping. One example is the enzyme AbaSI, which recognizes glucosylated 5hmC with high specificity when compared to 5mC or C, and cleaves the target sequences at a narrow range of distances away from the recognized 5hmC ^55^. The rare DNA bases 5fC and 5caC, which are thought to be intermediate bases in DNA demethylation reactions initiated by the TET family of 5mC oxidase proteins, can be removed from DNA by thymine DNA glycosylase (TDG) ^53,56,57^. Therefore, TDG could be used to map 5fC and 5caC at base resolution using circle-damage-seq.

The advantages of circle-damage-seq over other DNA damage mapping techniques are several-fold. (i) The single incision point of the modified base and the divergent reads emanating from this break position give a clear indication where the damage is located. (ii) Background signals are minimized by removal of DNA molecules that did not undergo circularization. (iii) Single strand breaks that also contribute to background are removed during circular ligation. (iv) The method represents adduct-specific sequencing because DNA circles or linear molecules without adducts are not sequenced, thus contributing to lower background and lower sequencing costs.

We have used circle-damage-seq for deriving information about the precise DNA sequence contexts of CPDs and etheno-dA adducts in exposed cells. This goal can easily be achieved with relatively low to moderate sequencing depth. However, for most DNA adducts it is still challenging to achieve a saturating sequencing density, meaning that all base positions of the genome are covered by enough reads to quantitate the base damage at each nucleotide position. Even when we used 0.8 billion reads for mapping of CPDs, we have not achieved this required depth. The reason is that dipyrimidines, the targets of CPD formation, occur on average at every 4^th^ base position on each DNA strand giving rise to approximately 800 million theoretical CPD positions per strand and haploid mammalian genome. It is likely that DNA sequencing costs will further decline in the future which will make the circle-damage-seq technology more generally affordable. When such high sequencing depth is achieved, it should also become feasible to follow precise DNA repair kinetics at every damaged base position of the genome over time and create high resolution DNA repair maps for different types of damaged bases.

Even with the current sequencing depth, we were able to clearly define the nearest neighbor sequence specificity of CPDs and observed that it is limited to a tetranucleotide context, which had not been appreciated previously. The formation of excited states in DNA by UV radiation is a key mechanism leading to photochemical base dimerization ^58^. These excited states can be delocalized over two or more stacked DNA bases and can be of either Frenkel exciton (ππ*) type, of a charge transfer type or represent a mix of both ^59–61^. The degree of sequence-specific base delocalization of excited states as well as the exact mechanisms how excited states undergo relaxation are still unclear, but our data suggest that delocalization does not extend beyond a tetranucleotide range.

Furthermore, we observe a pronounced reduction of CPD formation near the TSS and promoter regions of many genes (Fig. 5). The lower UV damage susceptibility of these critical gene control regions may be related to at least three phenomena: (i) DNA sequences at the TSS and within promoters are often G+C-rich, which provides a lower frequency of 5’TT or 5’TC sequences compared to the rest of the genome. These sequences are also generally unmethylated and less prone to CPD formation at 5-methylcytosine bases ^34,62^ (ii) Although a few transcription factors can enhance CPD formation when bound to DNA ^14,25,35,38^, most of these factors actually protect the DNA from forming pyrimidine dimers, presumably because the bound proteins provide more rigidity to their target DNA thus preventing the needed flexibility to form pyrimidine dimers ^14,25,26,36,37^. Perhaps multi-protein complexes consisting of general transcription factors involved in transcription initiation will also contribute to protection of DNA from UV damage. (iii) Sequences near the TSS are some of the most rapidly repaired regions of the genome ^63^. Although we isolated DNA immediately after irradiation, some repair may have already taken place. The exclusion of CPDs from these critical gene control regions should permit the maintenance of functional gene expression programs in UV-exposed cells. We also observed a distinct enhancement of CPD formation at the transcription end sites (Fig. 5). This enhanced damage formation, most likely due to an A/T-rich sequence context near polyadenylation sites, might represent a barrier to transcript processing and could impact cell survival if the damage is not effectively repaired. This phenomenon may contribute to UVB-induced cytotoxicity.

The circle-damage-seq method will allow the definition of precise sequence contexts for DNA adduct formation, even at moderate sequencing depth, as we have shown here for CPDs and etheno-dA adducts. The comparison of these sequence contexts, once defined for a variety of mutagenic DNA adducts, with cancer genome sequencing data from hundreds or thousands of patients should be useful to identify candidate agents that may have an etiological link to specific human malignancies.

## METHODS

### Cell culture

Mouse embryonic fibroblast (MEF) cells from C57BL/6 mouse embryos were obtained from ATCC (Cat. SCRC-1008), grown in DMEM high glucose (Invitrogen; Carlsbad, CA) containing 15% fetal bovine serum and used at passage number <5. THLE-2 cells derived from primary human liver epithelial cells were obtained from ATCC (Cat. CRL-2706) and grown in BEGM medium (Lonza; Walkersville, MD) supplemented with 10% fetal bovine serum, 5 ng/ml EGF and 70 ng/ml phosphoethanolamine on the culture plates precoated with a mixture of 0.01 mg/ml fibronectin, 0.03 mg/ml collagen type I and 0.01 mg/ml BSA.

### UVB irradiation, 4-HNE treatment and genomic DNA isolation

To irradiate cells with ultraviolet B (UVB) light, MEF cells at 80-90% confluence on 10 cm culture plates were washed with 1x PBS and irradiated in PBS with various doses (600-1500 J/m^2^) of UVB. The UVB lamp (ThermoFisher Scientific, UVP 3UV lamp) has a peak spectral emission at 302 nm. The UVB dose was determined using a UVX radiometer with a UVB probe (Ultraviolet Products; Upland, CA). To induce 1,*N*^6^-etheno-deoxyadenosine (etheno-dA) adducts in human cells, the THLE-2 liver cells at 80-90% confluence on 10 cm culture plates were washed with 1x PBS and treated with 40 μM of 4-HNE (Cayman; Ann Arbor, MI) in PBS for 10 min at 37°C.

After the irradiation or chemical treatment, the cells were immediately trypsinized and collected, and then genomic DNA was isolated using Quick-DNA Miniprep Plus Kit (Zymo Research; Irvine, CA) according to the manufacturer’s instruction manual.

### Circle-damage-seq mapping of CPDs

#### DNA preparation and circularization

To prepare fragmented genomic DNA, the DNA from UVB-irradiated cells was sheared to an average length of 300 bp by sonication with a Covaris E220 sonicator (Covaris; Woburn, MA) under the following conditions: peak incident power (W), 140; duty factor, 10%; cycles per burst, 200 times, 80 seconds, and then the fragmented DNAs were purified with DNA Clean & Concentrator-5 Kit (Zymo Research, DCC-5) according to the manufacturer’s protocol. DNA was eluted in 42 μl of 10 mM Tris-HCl pH 7.5. Next, the sonicated DNA was end-repaired to prepare blunt-ended DNA. One microgram of the sonicated DNA was incubated in 1x T4 DNA ligase buffer (NEB; Ipswich, MA) with components; 50 mM Tris-HCl, pH 7.5, 10 mM MgCl_2_, 1 mM ATP and 10 mM DTT and with 1 μl (2 U/μl) of RNase H (NEB), 1 μl (3 U/μl) of T4 DNA polymerase (NEB), 1 μl (10 U/μl) of T4 polynucleotide kinase (NEB) and 1 μl of 10 mM dNTPs in a final volume of 50 μl at 24°C for 30 min, and then the reaction was heat-inactivated by further incubation at 75°C for 20 min. The DNA was purified using DCC-5 Kit (Zymo Research) and eluted in 100 μl of 10 mM Tris-HCl, pH 7.5. The purified DNA was quantitated using Nanodrop (Thermo Fisher Scientific).

To circularize the DNA, 1 μg of the blunt-ended DNA was further incubated in 1x T4 DNA ligase buffer (NEB) in 1x T4 DNA ligase buffer (NEB) with 4 μl (400 U/μl) of T4 DNA ligase (NEB) in a final volume of 200 μl at 16°C overnight. Then, the reaction mixture was cleaned up with DCC-5 kit and eluted in 42 μl of 10 mM Tris-HCl, pH 7.5. To remove non-circularized linear DNA from the circularized DNA pool, the ligase-treated DNA was incubated with 2 μl (10 U/μl) of Plasmid-Safe ATP-Dependent DNase (Lucigen; Middleton, WI) in 1x reaction buffer (33 mM Tris-acetate, pH 7.5, 66 mM potassium acetate, 10 mM magnesium acetate, 0.5 mM DTT and 1 mM ATP) in a final volume of 50 μl at 37°C for 30 min and then at 70°C for 20 min to stop the reaction. Then, the DNA was cleaned up with 90 μl (1.8x) of AMPure XP beads (Beckman Coulter; Indianapolis, IN) and eluted in 35 μl of 10 mM Tris-HCl, pH 8.0.

#### Cleavage of DNA at CPD sites

To generate double-strand breaks at CPD sites, the circularized DNA was treated with the following DNA repair enzymes. First, to specifically incise DNA at CPD positions, the circularized DNA was incubated in 1x NEBuffer 4 reaction buffer (NEB) (50 mM potassium acetate, 20 mM Tris-acetate, pH 7.9, 10 mM magnesium acetate, 1 mM DTT) and 0.4 μl of 20 mg/ml BSA with 1 μl (10 U/μl) of T4 endonuclease V (NEB, T4-PDG) and 1 μl (10 U/μl) of APE1 (NEB) in a final volume of 40 μl at 37°C for 20 min. Then, the DNA was cleaned up with 72 μl (1.8x) of AMPure XP beads and eluted in 25 μl of 10 mM Tris-HCl, pH 7.5. To revert the dimerized pyrimidines remaining after T4-PDG incision, the DNA from above was treated with 3 μl (0.25 μg/μl) of *E. coli* phrB photolyase (Novus Biologicals; Centennial, CO) in 1x reaction buffer (50 mM Tris-HCl, pH 7.0, 50 mM NaCl and 10 mM DTT) in a final volume of 50 μl. The reaction tube was placed at a distance of 15 cm under a UVA lamp (ThermoFisher Scientific) that has a peak spectral emission at 365 nm and incubated under UVA illumination for 90 min at room temperature. Then, the DNA was cleaned up with 90 μl (1.8x) of AMPure XP beads and eluted in 32 μl of 10 mM Tris-HCl, pH 8.0. To cleave the opposite strand of the nicked DNA, the DNA from above was incubated with 1 μl (5 U/μl) of single-strand specific S1 nuclease (ThermoFisher Scientific) in 1x reaction buffer (40 mM sodium acetate, pH 4.5, 300 mM NaCl and 2 mM ZnSO_4_) in a final volume of 40 μl for 4 min at room temperature. We stopped the reaction by adding 2 μl of 0.5 M EDTA and 1 μl of 1M Tris-HCl, pH 8.0 to the reaction mixture and further incubated the mixture for 10 min at 70°C. The DNA sample was cleaned up with 72 μl (1.8x) of AMPure XP beads and eluted in 48 μl of 10 mM Tris-HCl, pH 8.0.

#### Adapter ligation

The double-strand cleaved DNA was subsequently A-tailed in 1x NEBNext dA-tailing reaction buffer (NEB) (10 mM Tris-HCl, pH 7.9, 10 mM MgCl_2_, 50 mM NaCl, 1 mM DTT) and 0.2 mM dATP, with 2 μl (5 U/μl) of Klenow fragment exo- (NEB) in a final volume of 55 μl by incubation at 37°C for 30 min, and then the reaction was heat-inactivated by further incubation at 70°C for 30 min. To perform adaptor ligation, 30 μl of NEBNext Ultra II Ligation Master Mix, 1 μl of NEBNext Ligation Enhancer (NEB), and 2 μl of 1.5 μM NEBNext adaptors for Illumina sequencing (NEB): 5’-phos-GATCGGAAGAGCAC-ACGTCTGAACTCCAGTC/ideoxyU/ACACTCTTTCCTACACGACGCTCTTCCGATC*T-3’ and 5’-phos-GATCGGAAGAGCACACGTCTGAACTCCAGTC/ideoxyU/AC-ACTCTTTCCTACACGACGCTCTTCCGATC*C-3’ (*; phosphorothioate linkage) were added to the dA-tailing reaction mixture in a total volume of 93 μl and incubated at 20°C for 15 min, and followed by incubation with 3 μl (1 U/μl) of USER enzyme (NEB) at 37°C for 20 min. Then the adaptor ligated circle-damage-seq library was cleaned up with 96 μl (1x) of AMPure XP beads and size-selected (300 to 1,000 bp) with 0.5 volumes of AMPure XP beads and eluted in 25 μl of 10 mM Tris-HCl, pH 8.0.

#### Library amplification and sequencing

To amplify the circle-damage-seq library by PCR, the eluted library was mixed with 25 μl of NEBNext Ultra II Q5 Master mix (2x), 2.5 μl of 10 μM NEBNext i5 and 2.5 μl of 10 μM NEBNext i7 primers (NEB) in a final volume of 50 μl. The amplification reaction mixture was incubated at 98°C for 30 sec, and then 12 cycles of PCR at 98°C for 10 sec and 65°C for 75 sec were performed, followed by a final extension step at 65°C for 5 min. The PCR product was purified with 50 μl (1x) of Ampure XP beads and eluted in 20 μl of 10 mM Tris-HCl, pH 8.0. The purified library was quantitated using a Qubit 3.0 fluorometer and ds DNA HS Assay kit (Thermo Fisher Scientific). The size distribution of the circle-damage-seq library was determined using Bioanalyzer (Agilent; Santa Clara, CA), and the circle-damage-seq library was quantified using KAPA Library Quantification kit (Kapa Biosystems) according to the manufacturer’s instruction. The circle-damage-seq library was sequenced as 150 bp paired-end sequencing runs on an Illumina Hiseq2500 platform to obtain between 10 million and 800 million reads. From the 800 million read sample, approximately 200 million paired aligned reads were retained specifically as divergent read pairs with single base gaps after removal of duplicates.

### Circle-damage-seq mapping of etheno-dA adducts

To prepare fragmented DNAs, NlaIII restriction enzyme (NEB) that leaves 5’CATG overhangs was used for digestion of the 4-HNE treated genomic DNA. We initially optimized the enzyme digestion to produce an average length of 300 bp, and 2 μg of genomic DNA was treated with 1 μl (5 U/μl) of NlaIII in a total volume of 50 μl at 37°C for 2 min. The sheared DNA was purified using DCC-5 Kit and eluted in 100 μl of 10 mM Tris-HCl, pH 7.5. The purified DNA was quantitated using Nanodrop.

To circularize the sheared DNA, 1 μg of NlaIII-digested, sticky-ended DNA was incubated in 1x T4 DNA ligase buffer with 4 μl (400 U/μl) of T4 DNA Ligase (NEB) in a final volume of 200 μl at 16°C overnight. Then, the reaction mixture was cleaned up with DCC-5 kit and eluted in 42 μl of 10 mM Tris-HCl, pH 7.5. To remove non-circularized linear DNA from the circularized DNA pool, the ligase treated DNA was incubated with 3 μl (10 U/μl) of Plasmid-Safe ATP-Dependent DNase in a final volume of 50 μl at 37°C for 30 min and then at 70°C for 20 min to stop the reaction. Then, the DNA was cleaned up with 90 μl (1.8x) of AMPure XP beads and eluted in 27 μl of 10 mM Tris-HCl, pH 8.0.

To generate double-strand breaks at etheno-dA adduct sites, the circularized DNA was initially treated with human alkyl adenine DNA glycosylase (hAAG, NEB) to cleave the *N*-glycosidic bond of etheno-dA bases. The circularized DNA was incubated in 1x ThermoPol Reaction buffer (provided by NEB for the hAAG reaction) with components; 20 mM Tris-HCl, pH 8.8, 10 mM (NH_4_)_2_SO_4_, 10 mM KCl, 2 mM MgSO_4_ and 0.1% Triton X-100, with 1 μl (10 U/μl) of hAAG in a final volume of 30 μl at 37°C for 30 min. Then, the DNA was cleaned up with 54 μl (1.8x) of AMPure XP beads and eluted in 35 μl of 10 mM Tris-HCl, pH 7.5. Subsequently, the hAAG treated DNA was incubated in 1x NEBuffer 4 reaction buffer with 1 μl (10 U/μl) of T4 PDG (AP lyase) and 1 μl (10 U/μl) of APE1 (to double-incise the AP sites) in a final volume of 40 μl at 37°C for 20 min, and the nicked DNA was cleaned up with 72 μl (1.8x) of AMPure XP beads and eluted in 32 μl of 10 mM Tris-HCl, pH 7.5. To cleave the opposite strand of the nicked DNA, the DNA was incubated with 1 μl (5 U/μl) of single-strand specific S1 nuclease in 1x reaction buffer (see above) in a final volume of 40 μl for 4 min at room temperature. Then, the reaction was stopped by adding 2 μl of 0.5 M EDTA and 1 μl of 1M Tris-HCl, pH 8.0 to the reaction mixture. We further incubated the mixture for 10 min at 70°C, followed by purification with 72 μl (1.8x) of AMPure XP beads. We eluted the DNA in 48 μl of 10 mM Tris-HCl, pH 8.0. circle-damage-seq library preparation and amplification by PCR were performed as described above using dA-tailing and ligation of the (5’-phos-CGGTGGACCGATGATC/ideoxyU/ATCGGTCCACCG*T-3’) hairpin linker. The circle-damage-seq library for etheno-dA mapping was sequenced using 75 bp paired-end reads on an Illumina NextSeq500 platform to obtain 15 million read.

### Circle-damage-seq mapping of DpnI cuts in the *E. coli* genome

As a pilot experiment, we initially performed the mapping of DpnI cut sites to optimize the circle-damage-seq method. To test the sensitivity of circle-damage-seq towards different percentages of DpnI-cleaved sites, we mixed *E. coli* DNAs from two different strains, dam(+) and dam(-) to prepare 0%, 2% and 10% of dam(+) DNA in a background of dam(-) input DNA. We subjected this mixture to the circle-damage-seq procedure as described above. Briefly, to prepare fragmented DNAs, RsaI and AluI restriction enzymes (NEB) that leave blunt ends were used to digest 1 μg of *E. coli* genomic DNA. The cleaved DNAs were subjected to A-tailing using NEBNext Ultra II End Prep kit (NEB) and followed by ligation of an annealed hairpin adaptor (5’-phos-CGGTGGACCGATGATC/ideoxyU/ATCGGTCCACCG*T-3’) using NEBNext Ultra II Ligation Master mix. The adaptor-ligated DNAs were treated with USER enzyme and circularized with T4 DNA ligase followed by treatment with Plasmid-Safe ATP-Dependent DNase. To generate double-strand breaks at DpnI cleavage sites, the circularized *E. coli* DNAs were incubated in 1x CutSmart buffer (NEB) with 1 μl (20 U/μl) of DpnI restriction enzyme (NEB) in a final volume of 20 μl at 37°C for 1 hour. Then, circle-damage-seq library preparation and amplification by PCR were performed as described above. The circle-damage-seq library was sequenced as 75 bp paired-end reads on an Illumina NextSeq500 platform to obtain ~15 million reads.

### Bioinformatics analysis

All libraries were hard-clipped from the 3’ end to 2×50 nucleotide length using *Trim_Galore* (https://www.bioinformatics.babraham.ac.uk/projects/trim_galore/). Reads were then adapter-trimmed and quality-trimmed (phred <20) in a single pass using *Trim_Galore* default parameters. Trimmed reads were aligned using *BWA-MEM* ^64^ with default parameters. Duplicate alignments were marked and removed with the ‘removeDups’ *SAMBLASTER* ^65^ option. *SAMtools* ^66^ was then used to remove reads with low mapping quality (MAPQ <20), and to sort reads by genomic coordinates.

Divergently aligned read pairs (pairs facing away from each other) with a single nucleotide gap between reads were selected by selecting SAM records with TLEN of 3. TLEN equals the distance between the mapped end of the template and the mapped start of the template ^66^. Records with TLEN=3 represent divergently aligned reads with 1 nucleotide between mates. Sequences for the 3 nucleotides representing the last nucleotide of each read and the gapped base were identified using *BEDtools* ^67^.

When treating cells with UVB to produce CPDs, if the gapped nucleotide was T or C in the (+) strand, the damage would have occurred on the (+) strand, whereas a gapped A or G in the (+) strand would indicate that the damage must have occurred on the (-) strand. For 4-HNE treatment producing etheno-dA adducts, if the gapped nucleotide was A in the (+) strand, the damage would have occurred on the (+) strand, whereas a gapped T would indicate that damage must have occurred on the paired A in the (-) strand. Gapped G or C found in libraries produced following 4-HNE treatments provide no inference about the strand in which damage occurred. As seen in Fig. S1, the DpnI-treatment protocol probes 6mA formed at 5’GATC sequences from dam(+) *E.coli* producing sequencing libraries with no gap between divergently aligned read pairs. As such, alignment records with TLEN=2 were selected.

Chromosome name, start position, stop position, and inferred strand (+/-) for all selected read pair loci were written to a BED file, sorted, and indexed with *tabix* ^68^ for downstream analyses. Logo plots were drawn using ggseqlogo ^69^.

## Data availability

All data are available from the GEO database (submitted on October 20, 2020 with immediate public access).

## Code availability

All code for this project is available at https://github.com/deanpettinga/CDseq.

## Acknowledgements

This work was supported by NIH grant CA228089 to GPP.

## Author contributions

S.G.J. and G.P.P. conceptualized and designed the project. S.G.J. and J.J. performed experiments. D.P., S.G.J. and G.P.P. performed data analysis. G.P. and S.G.J. wrote the manuscript. All authors commented on the manuscript.

## Supplementary Figures

**Figure S1:**
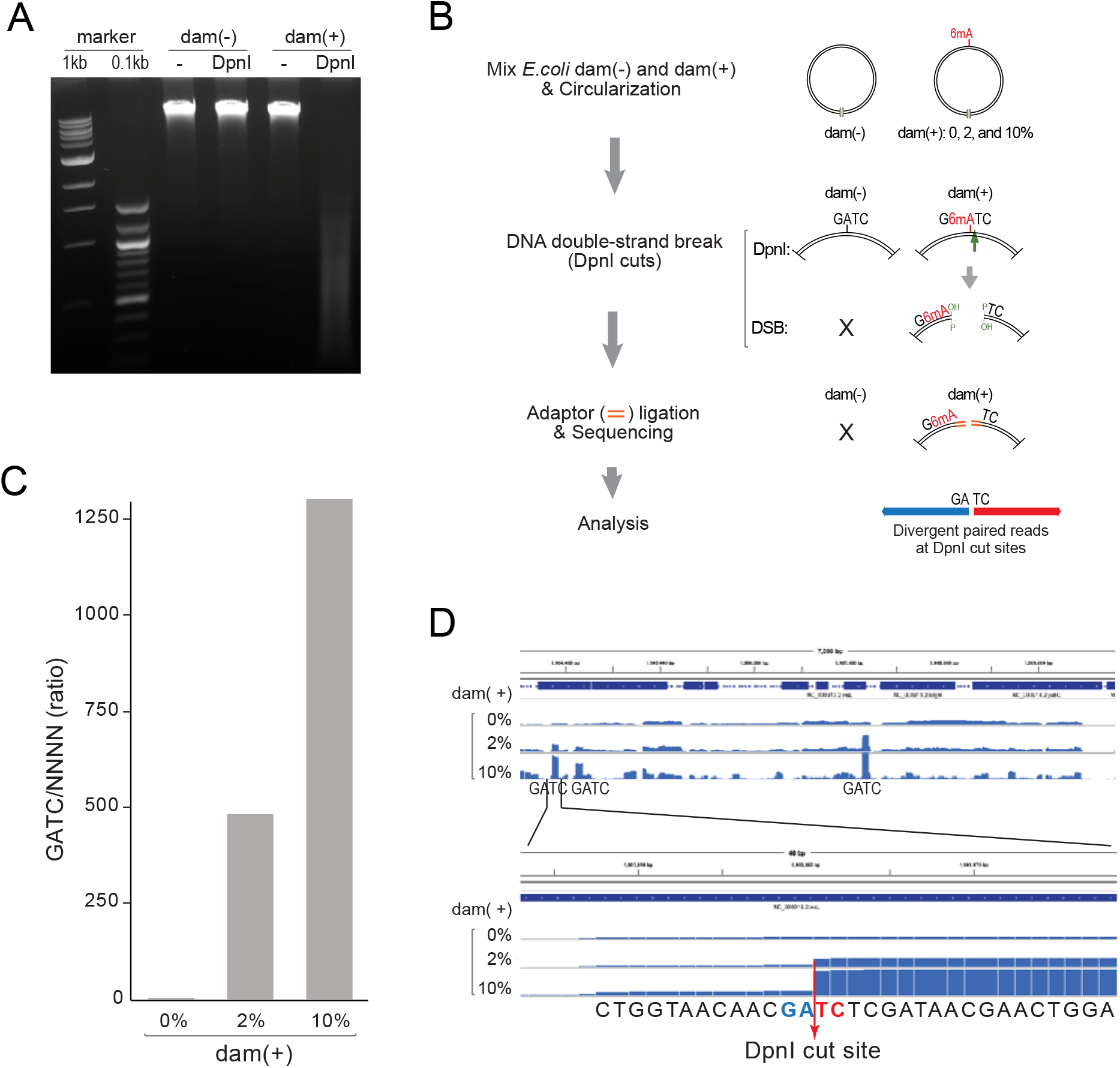
Mapping of N6-methyladenine at GATC sites. **A.** The agarose gel shows DpnI cleavage of DNA from dam(+) *E. coli* cells but not from dam(-) cells. **B.** Outline of the method to map DpnI cleavage sites. After circularization, DNA is cleaved at methylated GATC sites using DpnI. Linkers are ligated and the opened molecules are sequenced. Divergent, paired reads indicate the positions of methylated adenines. **C.** Ratio of cleaved 5’GATC sites relative to 5’NNNN sequences displayed as paired, divergent reads. **D.** Example of read stacks at methylated GATC sequences.

**Figure S2:**
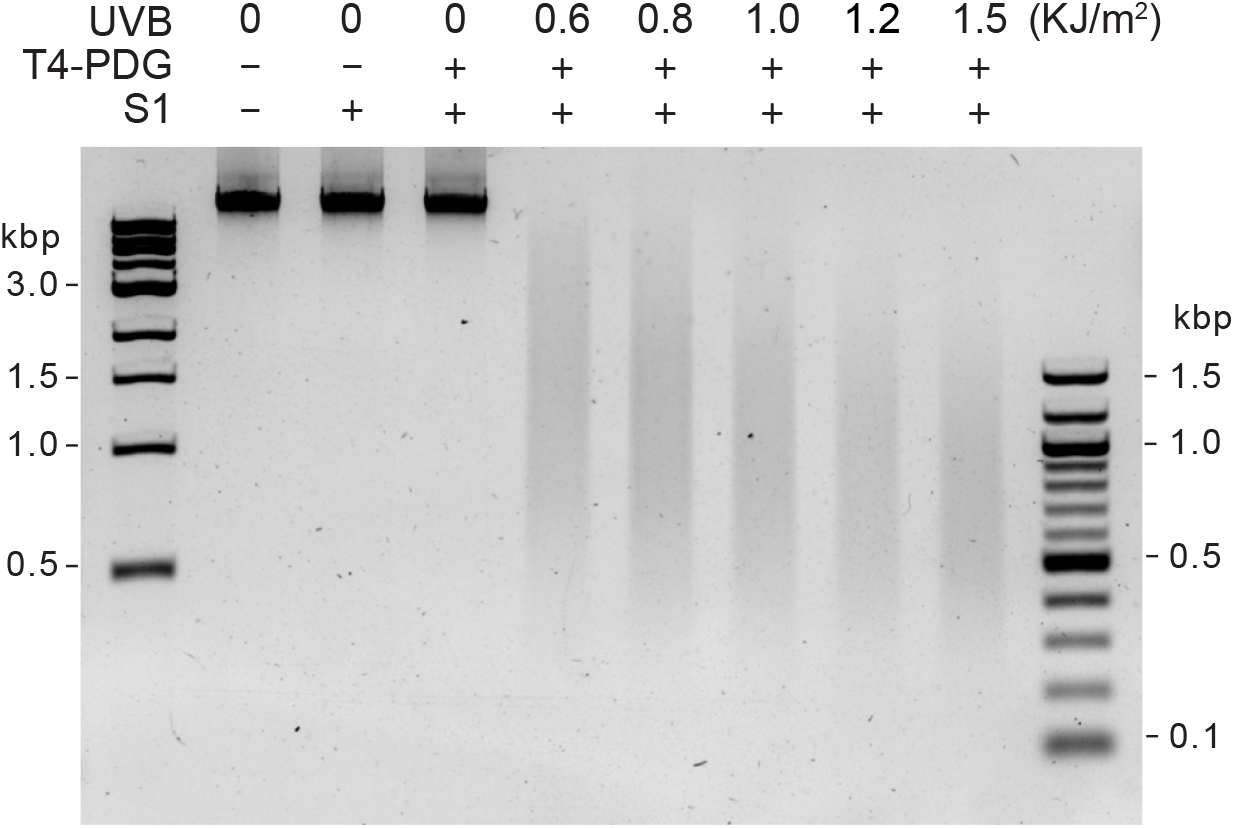
Double-strand cleavage at UVB-induced CPDs. Cells were irradiated with different doses of UV from a UVB light source. DNA was isolated and cleaved with the indicated enzymes. The cleaved DNA molecules were separated on a non-denaturing agarose gel. Double cleavage with T4 endonuclease V and S1 nuclease leads to dose-dependent double-strand break formation.

**Figure S3:**
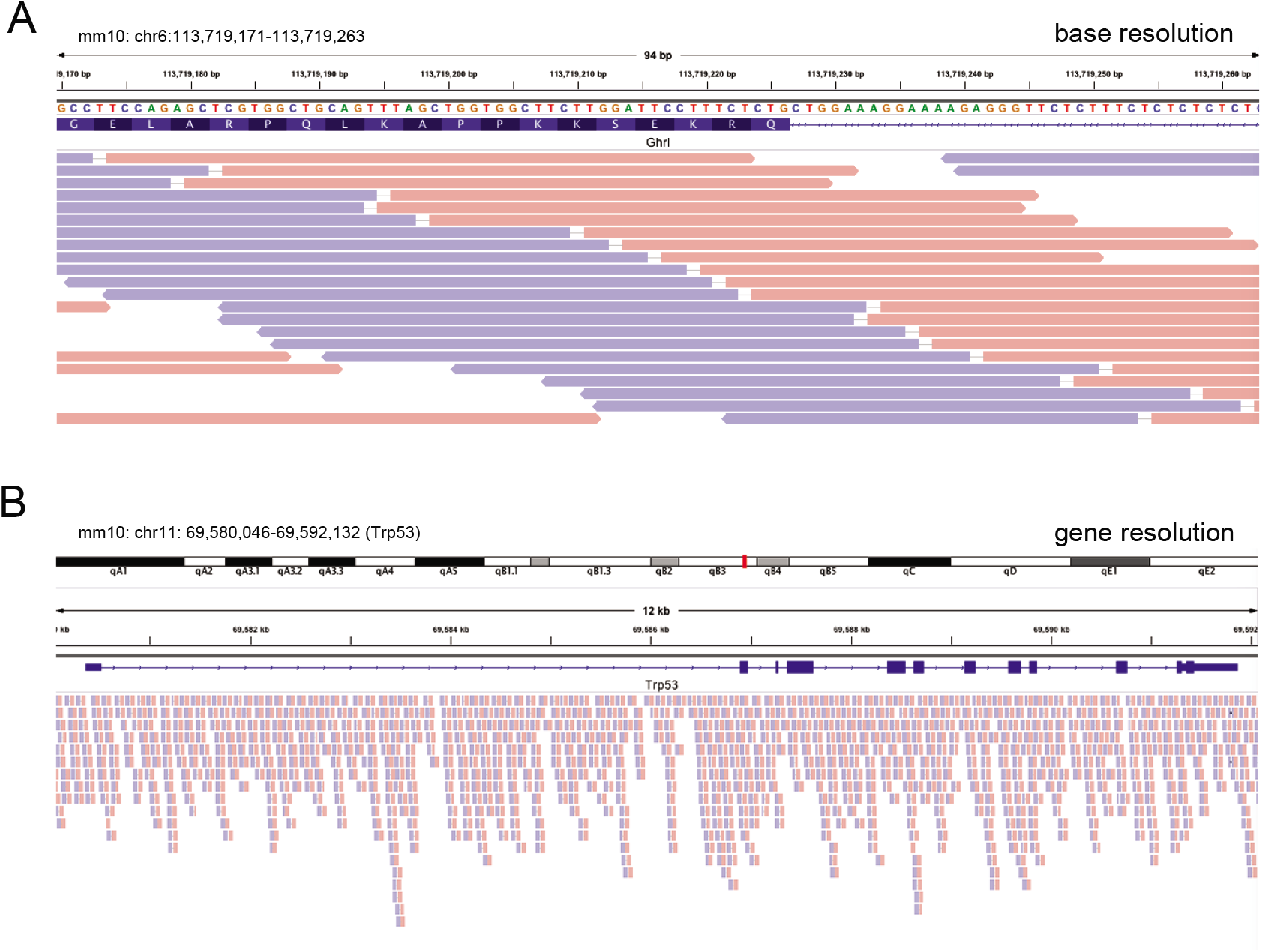
Example of CPD mapping by circle-damage-seq. **A.** Divergent reads (shown in red and blue colors for the forward and reverse reads) with single nucleotide gaps indicate the positions of UVB-induced CPDs. **B.** Lower resolution view of divergent reads from UVB-treated cells.

**Figure S4:**
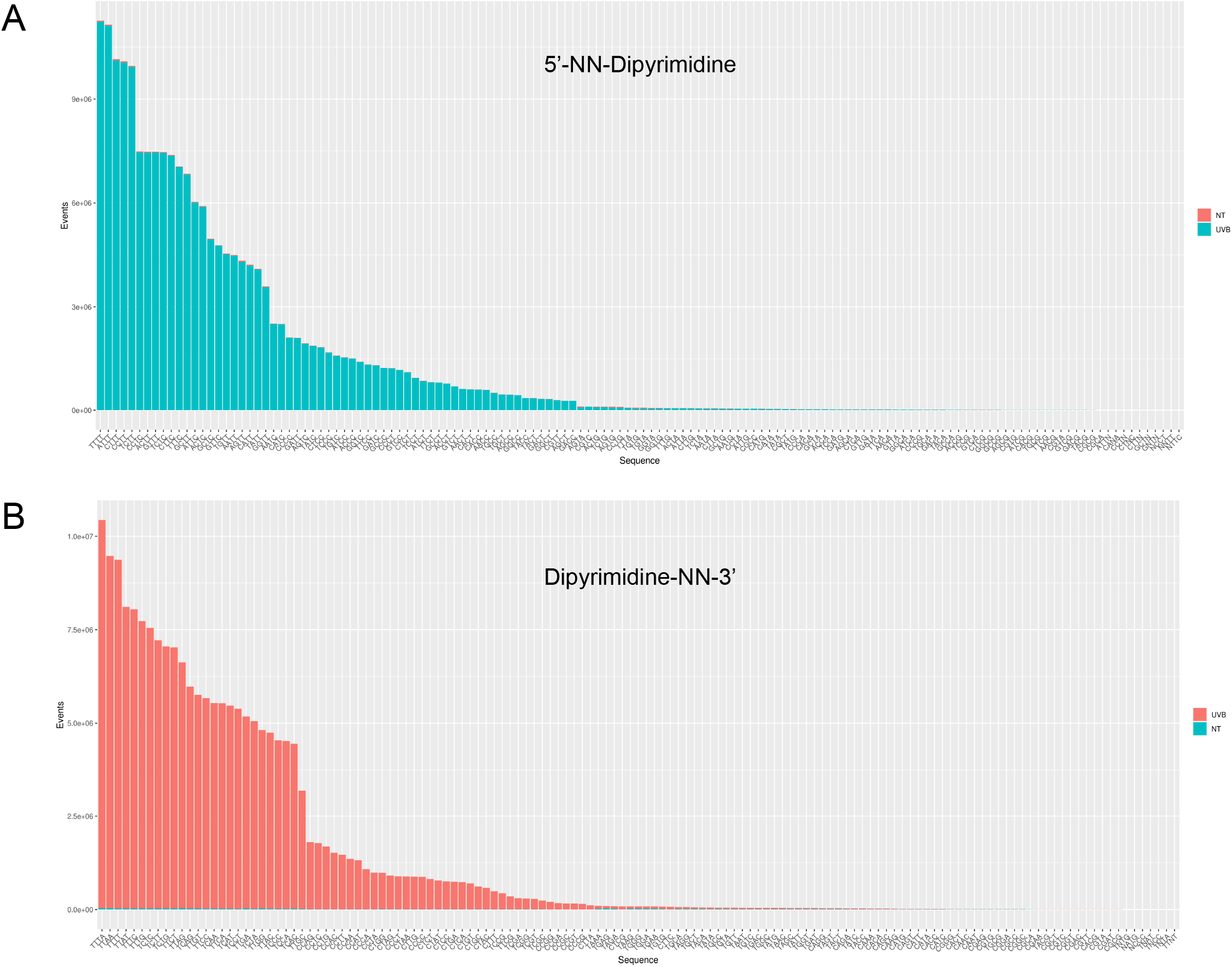
UVB-induced CPD tetranucleotide sequence specificity. **A.** Analysis of the two bases 5’ to a CPD. The sequence context is 5’NNY<>Y. **B.** Analysis of the two bases 3’ to a CPD. The sequence context is 5’Y<>YNN.

**Figure S5:**
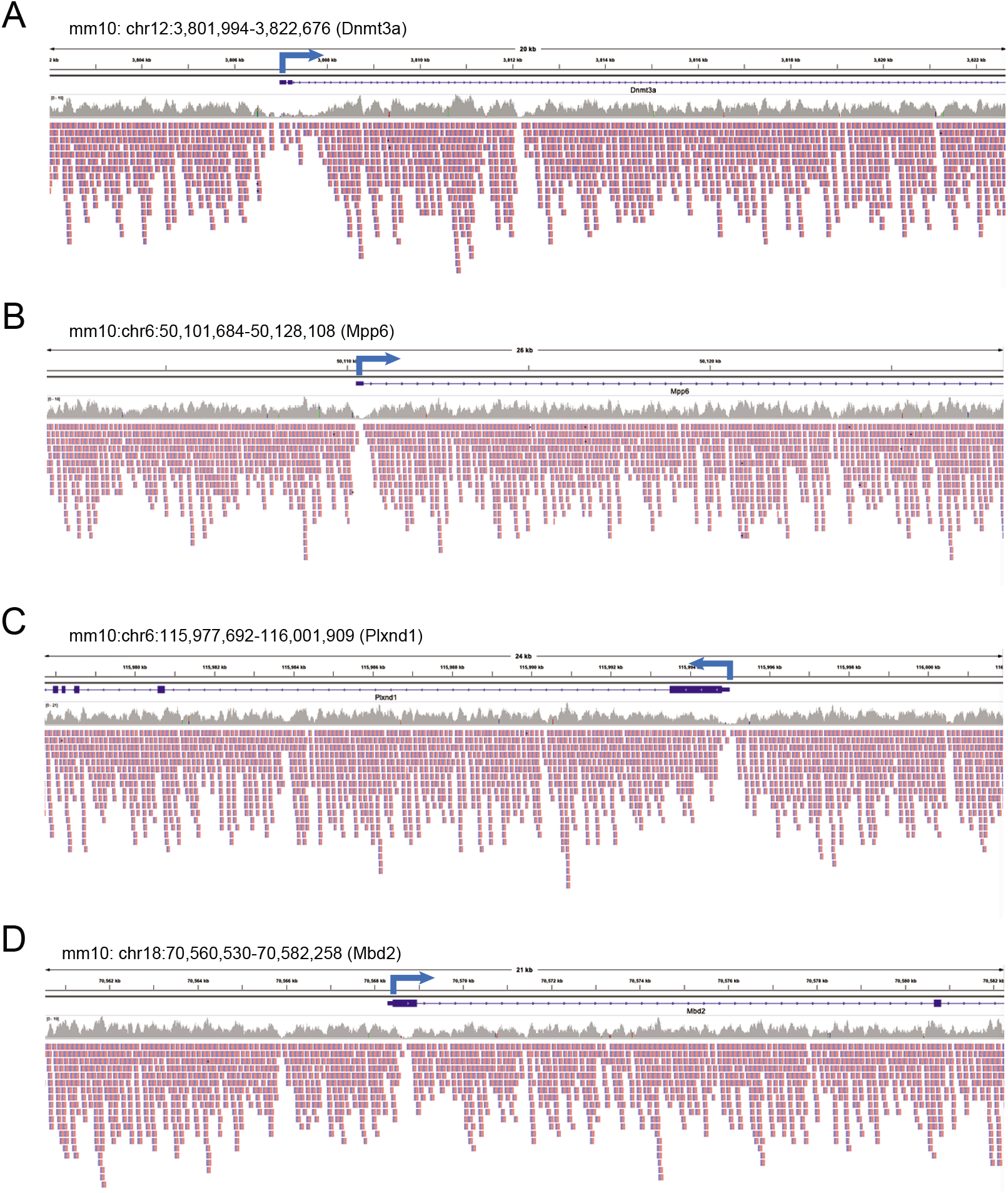
Reduced CPD levels at transcription start sites. These genes with G+C-rich promoters show the typical low CPD frequencies near the transcription start sites (TSS) (blue arrows indicate the direction of transcription). **A.** *Dnmt3a*. **B.** *Mpp6*. **C.** *Plxnd1*. **D.** *Mbd2*.

**Figure S6:**
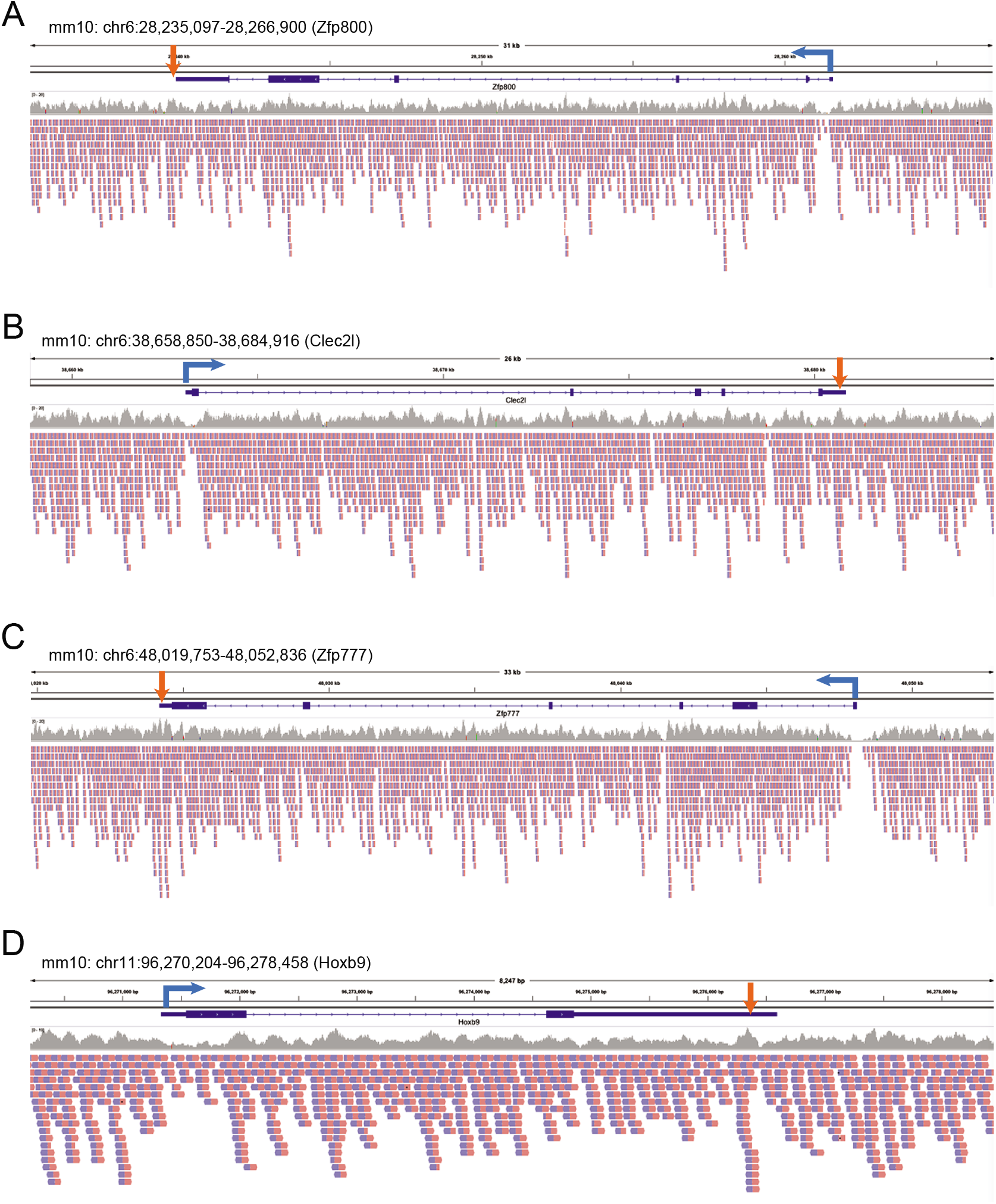
High CPD levels near transcription end sites. These examples show an increased level of CPDs near transcription end sites (TES) (red vertical arrows). **A.** *Zfp800*. **B.** *Clec2l*. **C.** *Zfp777*. **D.** *Hoxb9*. The low levels of CPDs near the TSS are also indicated (blue arrows).

